# Characterization and Validation of a Novel Group of Type V, Class 2 Nucleases for *in vivo* Genome Editing

**DOI:** 10.1101/192799

**Authors:** Matthew B. Begemann, Benjamin N. Gray, Emma January, Anna Singer, Dylan C. Kesler, Yonghua He, Haijun Liu, Hongjie Guo, Alex Jordan, Thomas P. Brutnell, Todd C. Mockler, Mohammed Oufattole

**Affiliations:** Benson Hill Biosystems, St. Louis, Missouri 63132, USA; Donald Danforth Plant Science Center, St. Louis, Missouri 63132, USA

## Abstract

CRISPR-based genome editing is an enabling technology with potential to dramatically transform multiple industries. Identification of additional editing tools will be imperative for broad adoption and application of this technology. A novel Type V, Class 2 CRISPR nuclease system was identified from *Microgenomates* and *Smithella* bacterial species (<u>C</u>RISPR from *<u>M</u>icrogenomates* and *<u>S</u>mithella*, Cms1). This system was shown to efficiently generate indel mutations in the major crop plant rice (*Oryza sativa*). Cms1 are distinct from other Type V nucleases, are smaller than most other CRISPR nucleases, do not require a tracrRNA, and have an AT-rich protospacer-adjacent motif site requirement. A total of four novel Cms1 nucleases across multiple bacterial species were shown to be functional in a eukaryotic system. This is a major expansion of the Type V CRISPR effector protein toolbox and increases the diversity of options available to researchers.

## Introduction

Genome editing, the ability to precisely alter DNA at a pre-determined location, has revolutionized research and development of novel applications. CRISPR (<u>c</u>lustered regularly interspaced <u>p</u>alindromic <u>r</u>epeats) nucleases that use DNA-RNA base pairing to provide site specificity have widely been adopted because of their low cost and ease of use relative to first-generation genome editing reagents that rely on DNA-protein interactions for site specificity. The first Class 2 CRISPR nuclease to be harnessed for genome editing was the Type II Cas9 nuclease from *Streptococcus pyogenes* (SpCas9)^1^. Following the demonstration of genome editing *in vitro* and in prokaryotes, SpCas9 was subsequently shown to function in eukaryotic cells^2^. Since the discovery and characterization of SpCas9, a number of additional diverse Type II Cas9 enzymes were discovered and harnessed for genome editing in various organisms^3-8^. The Cpf1 CRISPR nuclease family was subsequently the next family of CRISPR nucleases to be discovered and successfully used for genome editing^9^. Whereas Cas9 and Cpf1 enzymes both produced site-specific double-stranded breaks (DSBs) to effect genome editing, the nucleases shared little sequence identity beyond the fact that both Cas9 and Cpf1 nucleases have RuvC domains, a difference that resulted in the classification of Cpf1 enzymes as Type V nucleases. Accordingly, substantial additional investment has been made in thoroughly characterizing these groups^10-13^.

Since the discovery of Cpf1 enzymes, analyses of genomic and metagenomic data have uncovered additional Type V enzymes^13, 14^. Some of the newly discovered enzymes and putative enzymes have been validated for DSB production in heterologous systems^13, 15^, while others were tentatively identified based on computational analyses and await biochemical characterization. These newly discovered nucleases are likely to differentiate themselves for genome editing applications based on their varied physical and biochemical properties. Nucleases with smaller size, higher efficiency, higher specificity, divergent protospacer-adjacent motif (PAM) site requirements, and varied kinetic properties will expand the toolbox for researchers endeavoring to use genome editing. Here, we describe the discovery and characterization of a novel group of Type V, Class 2 enzymes that we term Cms1 (<u>C</u>RISPR from *<u>M</u>icrogenomates* and *<u>S</u>mithella*); these nucleases may also be classified as cas12e nucleases according to previous classifications^16^. These nucleases are smaller than most CRISPR nucleases, have a simple single RNA component, and have an AT rich PAM site requirement.

## Results

### Identification of a new group of Type V Nucleases

The initial description of Cpf1 nucleases included a phylogenetic tree showing an out group of four enzymes derived from *Smithella* sp. SCADC (SmCms1), *Smithella* sp. K08D17 (Sm2Cms1), *Microgenomates* sp. (MiCms1), and *Sulfuricurvum* sp. PC08-66 (SuCms1) that appeared to be an outgroup and did not have strong sequence homology to the characterized Cpf1 nucleases^9^. Further examination of these sequences showed that they were significantly divergent from Cpf1 amino acid sequences and likely represented a new group of Type V nucleases, which we named Cms1 (Figure 1A). Subsequently, we identified an additional enzyme from *Omnitrophica* bacterium (ObCms1) that has strong homology to this group of nucleases. Table 1 summarizes proteins described herein and provides the GenBank accession to the genomic contigs these Cms1 gene sequences were derived from.

**Figure 1:**
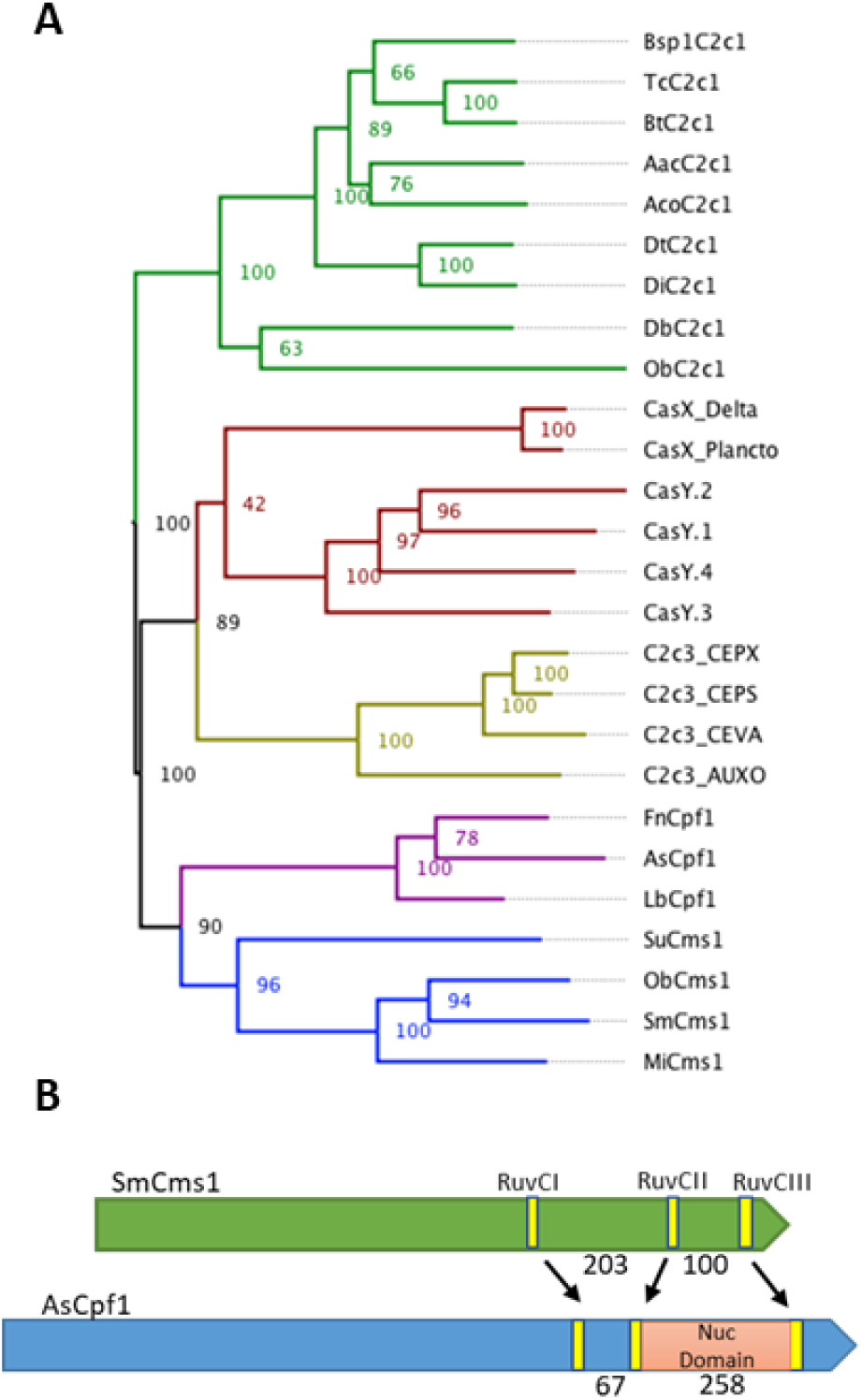
The Cms1 nuclease is a structurally different Type V CRISPR nuclease protein. **(A)** Midpoint-rooted maximum likelihood phylogenetic tree of Type V CRISPR nucleases. Numbers at nodes indicate bootstrap value support. **(B)** Comparison of RuvC domain spacing and presence/absence of Nuc domain between SmCms1 and AsCpf1 nucleases. Values are the number of amino acids between the annotated RuvC domain active sites.

**Table 1.** Description of Cms1 Nucleases.

| <u>Cms1 Protein</u> | <u>Organism</u> | <u>GenBank Protein Accession</u> | <u>GenBank DNA Accession</u> | <u># spacers present</u> |
| --- | --- | --- | --- | --- |
| SmCms1 | *Smithella* sp. SCADC | KFO67988 | JQDQ01000121 | 39 |
| MiCms1 | *Microgenomates* sp. | KKQ38176 | LBTJ01000016 | 8 |
| ObCms1 | *Omnitrophica* bacterium | OGX23684 | MHGE01000059 | 10 |
| SuCms1 | *Sulfuricurvum* sp. PC08-66 | KIM12007 | JQIT01000003 | 6 |

We identified substantive differences between Cpf1 and Cms1 nucleases using RuvC-anchored amino acid alignments of nucleases (Supplementary Figure 1). The RuvC domains contain the DNAase active site residues in CRISPR nucleases^17, 18^. The Nuc domain, located between the RuvCII and RuvCIII domains in Cpf1 proteins^18^, is absent in Cms1 proteins. In contrast, Cms1 proteins contain a large insertion between the RuvCI and RuvCII domains relative to Cpf1 proteins. BLAST and HHPred analyses of this domain failed to clearly identify the source or putative function of this domain. In addition, other smaller differences included multiple blocks of conserved or semi-conserved sequences found only in Cms1, whereas other conserved sequences were found only in Cpf1. Supplementary Table 1 summarizes the results of BLASTP and CLUSTALO alignments of Cms1 proteins with the Cpf1 proteins from *Francisella novicida* (FnCpf1) and from *Acidaminococcus* sp. (AsCpf1), with just 10-15% sequence identity shared between Cms1 nucleases and these Cpf1 nucleases. The relatively small sizes of the Cms1 proteins were not the cause of the low sequence coverage and identity values, as similar alignments between FnCpf1, AsCpf1, and a 1,154 amino acid Cpf1 protein derived from *Proteocatella sphenisci* resulted in 99% sequence coverage and 31-37% overall sequence identity, similar to the 100% coverage and 35% sequence identity shared between FnCpf1 and AsCpf1 (data not shown).

As in other previously reported Type V nucleases, Cms1 nucleases contain RuvC domains near their C-terminus that are divided into three subdomains. These RuvC domains and the anticipated active site residues were identified based on HMM analyses and sequence alignments with previously described Type V nucleases. Supplementary Table 1 shows the amino acid sequences surrounding the RuvCI, RuvCII, and RuvCIII active sites in the Cms1 nucleases described here and in previously described Type V nucleases. Specifically, the identified Cms1 nucleases do not contain the ADANG motif found in the RuvCIII domain of the listed Cpf1 nucleases and strongly conserved among other putative Cpf1 nucleases (Supplementary Table 2). Whereas the amino acid sequences of the Cms1 nucleases are most closely related to Cpf1, the spacing of the RuvC domains in the C-terminus is quite different, resulting from the presence of an unknown domain between RuvCI and RuvCII in Cms1 proteins and the presence of the Nuc domain between RuvCII and RuvCIII domains in Cpf1 proteins (Figure 1B and Table 2). These differences in sequence identify and structural orientation suggest a difference in overall structure-function.

Analysis of these bacterial genomic regions reveals that each identified gene was close to a CRISPR repeat region varying from 6-39 direct repeats (Figure 2A). Except for the nuclease from *Sulfuricurvum* sp. PC08-66, each Cms1 nuclease is also adjacent to known genes involved in CRISPR biology (Cas4, Cas1, and Cas2). Interestingly, the *Smithella* and *Microgenomates* Cms1 genes are adjacent to annotated Cpf1 nuclease genes. In addition, the *Smithella* CRISPR region has a second repeat region downstream with divergent direct repeat sequences, suggesting the presence of two divergent CRISPR systems at this locus (Supplementary Figure 2). While it is possible that the SmCms1 and MiCms1 genes could have resulted from duplication of the Cpf1 genes present in these genomes, BLASTN alignments showed extremely low sequence identity between Cpf1 and Cms1 genes, with only 6% coverage between the SmCms1 and SmCpf1 genes, and 6% coverage between the MiCms1 and MiCpf1 genes. Similarly, BLASTP alignments of the Cpf1 and Cms1 proteins encoded by these genes showed low coverage and low sequence conservation, similar to alignments between these Cms1 proteins and FnCpf1 and AsCpf1 proteins. Thus, colocalization of these genes within the genome may reflect similar functions in defense rather than a common origin.

**Table 2:** Spacing of RuvC domains in Cms1 and Cpf1.

| <u>Protein</u> | <u>RuvCI-RuvCII spacing (# amino acids)</u> | <u>RuvCII-RuvCIII spacing (# amino acids)</u> |
| --- | --- | --- |
| AsCpf1 | 84 | 269 |
| LbCpf1 | 92 | 254 |
| SmCms1 | 220 | 122 |
| ObCms1 | 211 | 113 |
| MiCms1 | 225 | 117 |
| SuCms1 | 214 | 149 |

**Figure 2:**
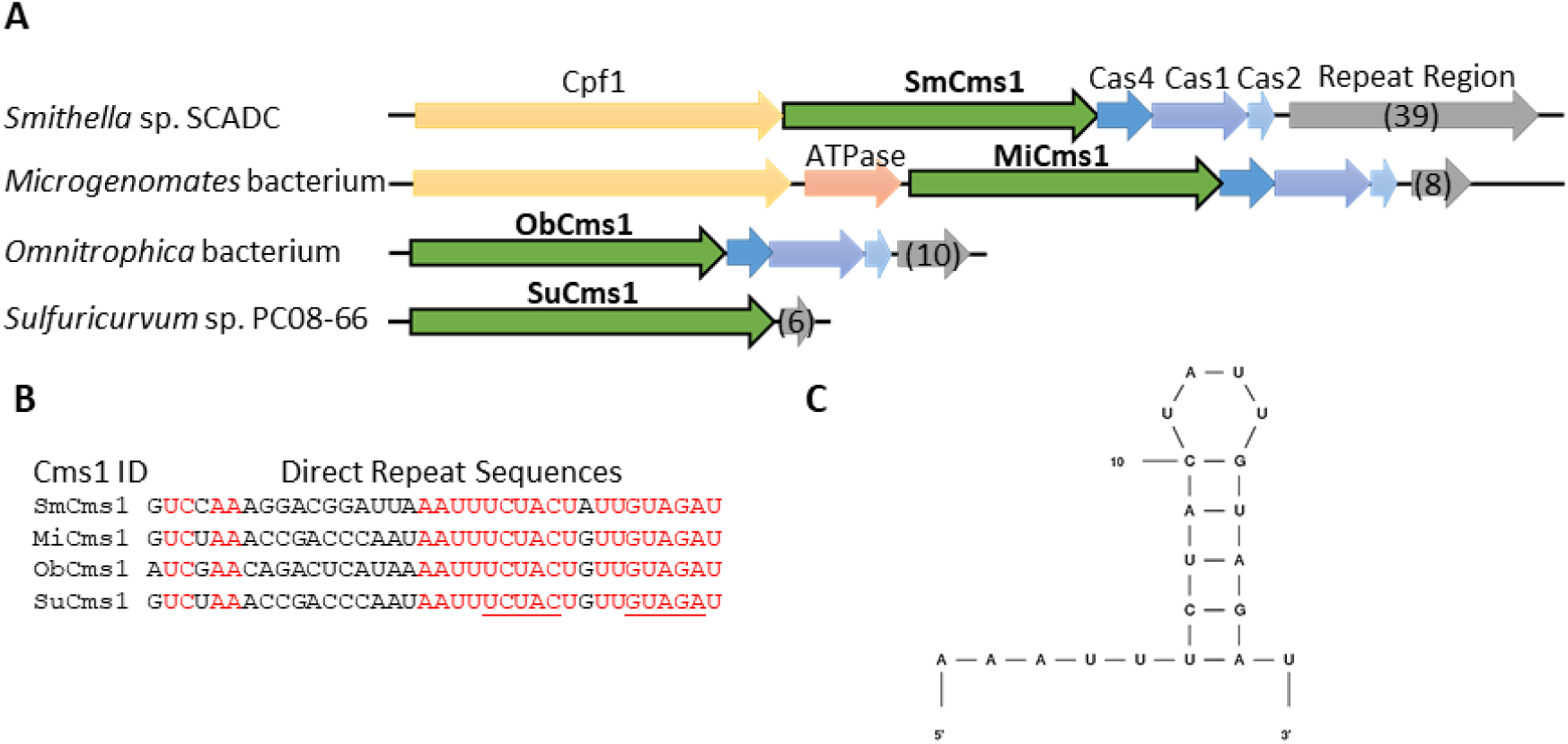
Cms1 nucleases are associated with CRISPR loci and have use a single RNA structure. **(A)** Orientation of Cms1 nucleases within bacterial genomes. Associated Cas proteins are color coded for clarity. The number of direct repeat sequences in each repeat region are in parentheses. **(B)** Alignment of the direct repeats sequences associated with each Cms1 nuclease. Conserved resides are highlighted in red. The predicted stem structure is underlined. **(C)** Predicted stem loop structure from SmCms1 direct repeat.

Each of these spacer sequences was flanked by a direct repeat sequence like those found in most Cpf1-encoding genomes^9^ (Figure 2B). Each of these direct repeats contained a conserved <u>TCTAC</u>TNTT<u>GTAGA</u> sequence near the 3’ end of the direct repeat, with the underlined bases predicted to form a hairpin structure (Figures 2C). Spacers in the CRISPR arrays associated with Sm, Mi, and ObCms1 genes had median sizes of 28, 29.5, and 27 bp, respectively, while spacers associated with SuCms1 were significantly larger with a median size of 32 bp. No tracrRNA sequence was identified upstream of the direct repeats in any of these genomic regions.

Additional evidence of differentiation was drawn from analyses of phylogenetic relationships. We identified subdomains based on catalytic amino acid residues, divided sequences into each subdomain, conducted multiple alignments on each, and then concatenated anchored multiple sequence alignments. A maximum likelihood tree was then derived from sequences described above, and bootstrapped to assess edge support of the tree (Figure 1A). Results confirmed previous classifications of previously described Cpf1, C2c1, C2c3, CasX, and CasY nucleases, and placed Cms1 nucleases on an entirely separate clade. Notably, Cms1 proteins clustered within the same clade, which was most closely related to Cpf1 nucleases. Bootstrap support for separation between the Cpf1 and Cms1 clades was high (90) and similar to differences between other well-recognized CRISPR families, further evidencing differentiation between Cms1 nucleases and other previously described proteins.

### *In planta* demonstration of SmCms1 as a functional nuclease

As part of a larger Type V CRISPR nuclease screening program to identify nucleases capable of efficient genome editing in plants, SmCms1 was tested for nuclease activity in rice (*Oryza sativa*). The experimental design for this screen is based on our original work with Cpf1 nucleases in rice callus material^19^. Briefly, rice callus material was bombarded with three separate plasmids (Supplementary Tables 3-4). The first plasmid contained the nuclease with an enhanced 35S promoter, the second plasmid contained the crRNA cassette with the rice U6 promoter and 24 bp target sequence for the rice CAO1 gene, and the third contained a repair template for the target site and included a hygromycin resistance marker with a maize (*Zea mays*) ubiquitin 1 promoter. A TTTC PAM site was used for this initial screen because it is compatible with most Type V nucleases. Due to the high throughput nature of this screen and the strong similarity between the direct repeat sequence of the SmCms1 CRISPR region and the FnCpf1 CRISPR region, the FnCpf1 crRNA hairpin structure was used. After four weeks of selection on hygromycin, the callus material was screened for the presence of indels via a T7EI assay. PCR products positive for the T7EI assay were sub-cloned and screened to validate the presence of edits. See Figure 3A for a schematic of the experimental design.

**Figure 3:**
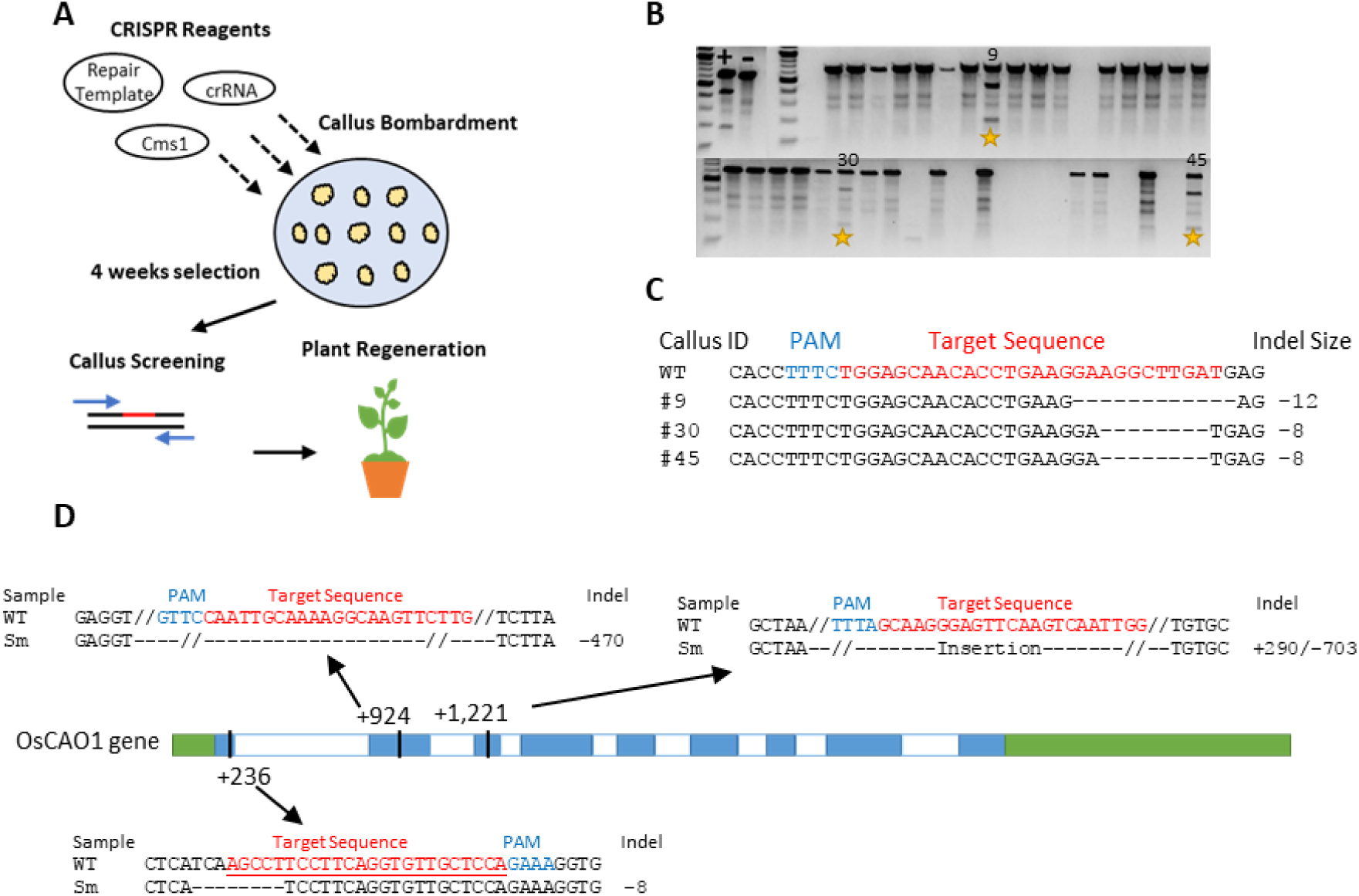
SmCms1 is a functional CRISPR nuclease with in vivo activity and an AT rich PAM site requirement. **(A)** Schematic of in planta nuclease screening system. **(B)** T7EI results from rice genome editing experiments with SmCms1. Positive and negative controls are shown with (+) and (-), respectively. Calli with sequence validated editing are highlighted with a yellow star and labelled. **(C)** Alignment of Sanger sequencing results from calli positive for indels. **(D)** Schematic of rice CAO1 gene and targets sites tested for nuclease activity. UTR regions shown in green, exons shown in blue, and introns are shown in white. Targets sites are labelled relative to the transcription start site of the gene.

Initially 48 hygromycin resistant callus were screened for the presence of indels by PCR amplification of the CAO1 target region and treatment with T7EI (Figure 3B). Clear indels were observed in callus samples 9, 30, and 45. The PCR products from these samples were sequenced and the resulting reads with indels were aligned to the wildtype sequence (Figure 3C). Indels of −8 and −12 were observed at the 3’ end of the target sequence. The median of the sequenced indels was 24 bp distal from the PAM site. This was the first demonstration of *in vivo* nuclease activity by a Cms1 nuclease. These results confirm that SmCms1 is a functional Type V CRISPR nuclease, does not require a tracrRNA, and can cut adjacent to a TTTC PAM site.

To date, attempts to develop an *in vitro* assay system for SmCms1 have been unsuccessful. Protein has been successfully purified from *E. coli*, but *in vitro* nuclease activity has yet to be detected in the buffer conditions and temperatures tested (See Supplementary Table 6). These *in vitro* assays are complicated by protein stability issues with SmCms1. Current work is being done to resolve this issue and develop a robust *in vitro* assay to further characterize this protein. While low solubility has been observed with other CRISPR nucleases^13^, it has limited our ability to generate a robust PAM site identification assay. In lieu of a functional *in vitro* assay, SmCms1 was tested at multiple sites in the rice CAO1 gene. These additional targets are outlined in Figure 3D along with the observed indel sequencing. SmCms1 was shown to generate indel mutations at three tested sites. Editing was detected at TTTA and GTTC PAM sites in addition to TTTC (Figure 3D). These data suggest that SmCms1 requires a TTN PAM site, but like many other Type V nucleases^13^,^20^, strongest activity appears to be at a TTTN PAM site.

**Figure 4:**
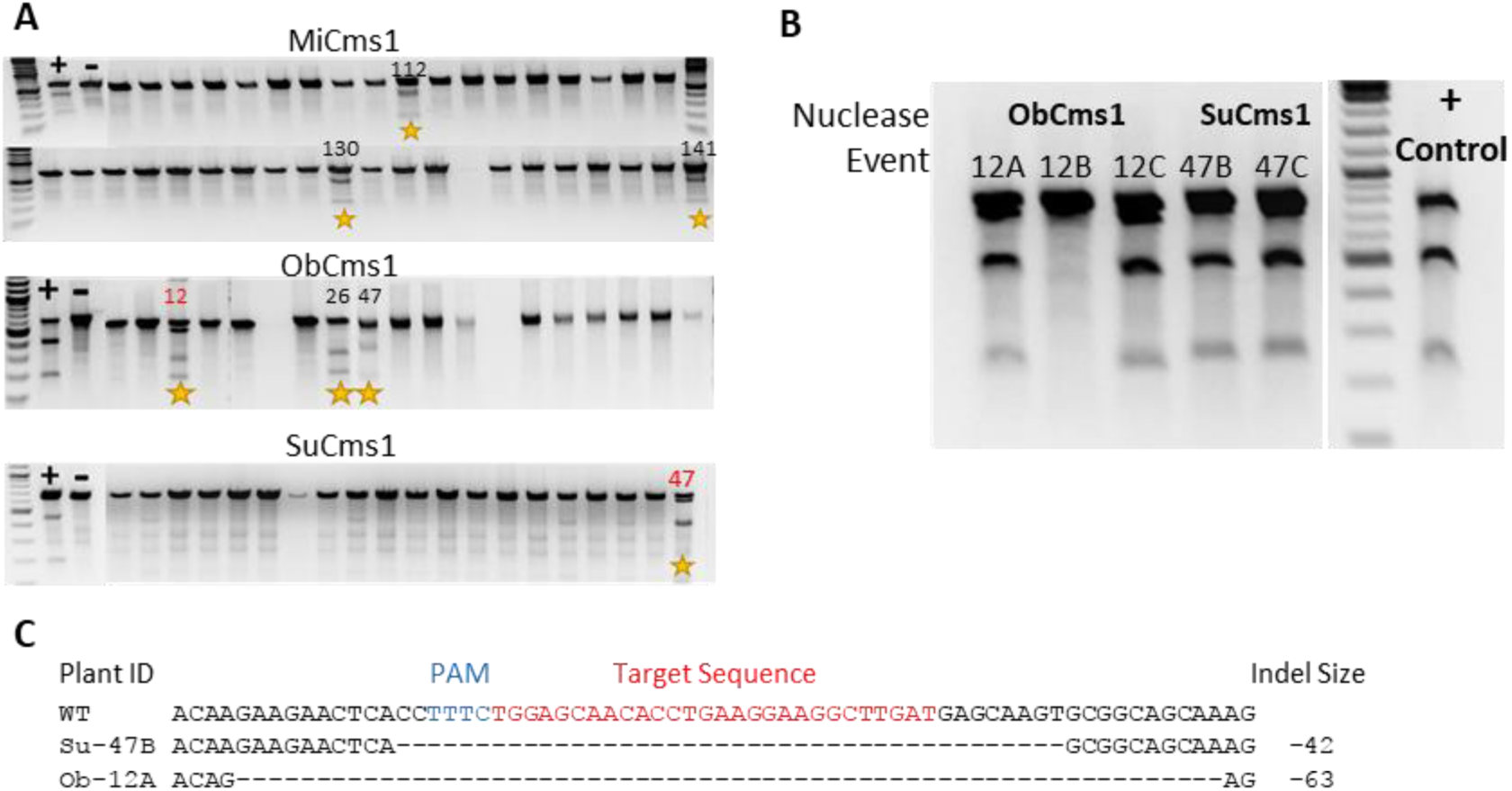
Cms1 nucleases from *Sulfuricurvum, Microgenomates*, and *Omnitrophica* are functional genome editing proteins *in vivo*. **(A)** T7EI results from rice genome editing experiments with SmCms1. Positive and negative controls are shown with (+) and (-), respectively. Calli with sequence validated editing are highlighted with a yellow star and labelled. **(B)** T7EI results from rice leaf tissues. Events from each nuclease are labelled above each lane. The respective nuclease is labelled above the plant events. Plants derived from the same piece of callus are labeled with number – letter nomenclature. **(C)** Sanger sequencing alignments of indels observed from rice callus and leaf tissue.

### Screening of additional Cms1 nuclease *in planta*

The additional Cms1 nucleases were run through the *in planta* rice screening system as previously described^19^. Indels were observed using the T7EI assay and validated via Sanger sequencing for three additional Cms1 nucleases: MiCms1, ObCms1, and SuCms1 (Figure 4A). As part of the screening system, it is possible to regenerate plants from sampled callus at a reasonable rate. Multiple T0 plants were regenerated from ObCms1 callus #12 and SuCms1 callus #47. These plants were screened for the presence of the edits that were observed in the callus stage. For each plant, a PCR product for the CAO1 target region was generated, run through a T7EI assay (Figure 4B), and sequenced (Figure 4C). Indels were observed for two of the plants derived from ObCms1 callus #12and mutations were observed in both sibling plants derived from SuCms1 callus #47. In all cases, the sequence of the indels in the plants matched the original sequencing from the callus material. All observed mutations appeared to be heterozygous due to the presence of a 1:1 sequence ratio of WT and mutant alleles for each plant. These collective data demonstrate the functionality of three additional Cms1 nucleases in an *in vivo* system.

## Discussion

The novel Type V, Class 2 Cms1 nucleases described and validated here for genome editing are among the smallest nucleases shown to be functional for eukaryotic genome editing to date. The small size of these nucleases, lack of any tracrRNA requirements, and *in planta* validation make Cms1 nucleases novel and valuable tools for eukaryotic genome editing (Table 3). Interestingly, all the Cms1 nucleases presented here demonstrated *in vivo* activity, which is strikingly different from reports of Cpf1 nucleases, where 12-25% of Cpf1 nucleases tested have supported eukaryotic genome editing^9, 21^. Expansion of the genome editing reagent tool box will continue to be important to broaden the application of this technology.

Based on their distant relationships to previously described Type V nucleases, presence of a RuvC domain, and absence of an HNH domain, Cms1 nucleases can be clearly classified as Type V CRISPR nucleases. Previous classifications of CRISPR nucleases have typically relied on analysis of not only the effector protein sequences themselves, but also on surrounding genomic contexts. This results in a classification scheme that is labor-intensive and has a degree of subjectivity associated with it ^11, 12, 14^. Under this paradigm, nucleases isolated from metagenomes or incomplete genomes where surrounding genomic context is unclear may not be unambiguously classified. Here, we describe a straightforward and reproducible method for classification of CRISPR nucleases that relies on first identifying subdomains based on catalytic amino acid residues, which then anchor subsequent multiple sequence alignments. This method was largely in agreement with previous classifications of the previously described Cpf1, C2c1, C2c3, CasX, and CasY nucleases, and clearly placed Cms1 nucleases on a separate clade with these nucleases being most closely related to Cpf1 nucleases. The absence of a Nuc domain and presence of an unknown domain located between the RuvCI and RuvCII domains in Cms1 nucleases (Figure 1B), as well as the very low sequence conservation between Cpf1 and Cms1 proteins (Supplementary Table 1) clearly supports the classification of these as a separate group of Type V nucleases (Figure 1A). The method described here should facilitate classification of CRISPR nucleases; these methods can be readily adapted to alignment of other proteins for phylogenetic analysis as well where conserved biologically active residues are known.

**Table 3.** Comparison of Type II and Type V CRISPR nuclease.

| Nuclease | CRISPR Type | Size (aa) | RNA Component | PAM Site |
| --- | --- | --- | --- | --- |
| SmCms1 | V | 1,065 | Single | TTN |
| SpCas9 | II | 1,368 | Dual | NGG |
| AsCpf1 | V | 1,307 | Single | TTTN |
| CasX | V | ~980 | Dual | TTCN |
| CasY | V | ~1,200 | Uncertain | TA |
| C2c1 | V | 1,130 | Dual | TTN |
| C2c3 | V | ~1,200 | Uncertain | Unknown |

Class 2 CRISPR systems described to date have typically included multiple Cas genes involved in spacer acquisition (i.e., Cas1, Cas2, and Cas4 genes) as well as an effector protein (e.g., Cas9 or Cpf1). Two of the Cms1 nuclease-encoding genes described here (SmCms1 and MiCms1) are found in CRISPR loci that also contain a Cpf1-encoding gene (Figure 2A). This organization, with a single CRISPR locus comprising more than one effector protein-encoding gene, has not been previously described to our knowledge. This unique organization raises questions about the biological function of the nucleases *in vivo* for bacterial immunity and other functions that may be regulated by these CRISPR systems.

The *in planta* genome editing mediated by multiple Cms1 nucleases at sequences downstream from AT-rich PAM sites demonstrates that PAM sites accessible by most Type V nucleases appear to be accessible by Cms1 nucleases as well (Table 3). The development of suitable *in vitro* assays for Cms1 nucleases will facilitate further elucidation of PAM site requirements for these nucleases as well as a deeper characterization of the underlying biochemistry. The identification and characterization of additional Cms1 nucleases will help with further characterization of this family of Type V nucleases. Additionally, given the quite distant amino acid similarity shared between Cms1 and other Type V nucleases, we anticipate that these nucleases will exhibit unique functionality when compared with other nucleases, and that the unique functionality can be harnessed for basic and applied genome editing activities.

## Methods

### *In silico* analyses

BLASTP and BLASTN searches and alignments were performed at NCBI (https://blast.ncbi.nlm.nih.gov/Blast.cgi) using default parameters. CLUSTALO protein alignments were performed at UniProt (http://www.uniprot.org/align) using default parameters^22^. HHPred protein analyses were performed using the Max Planck Institute for Developmental Biology Bioinformatics Toolkit (https://toolkit.tuebingen.mpg.de/hhpred), using default parameters.

The sequences of the Type V nucleases used for the analyses described here were obtained from Genbank entries listed in a previous publication describing CasX and CasY^13^ and from supplementary information provided in a publication describing C2c1 and C2c3^23^, and from Genbank entries listed in a previous publication describing Cpf1^9^.

Genome sequences were obtained from the National Center for Biotechnology Information (NCBI) in August 2017. Active site residues for RuvCI, RuvCII, and RuvCIII were identified in each of 22 Type V sequences and used to define domain boundaries. Sequences were then stratified into each of four domains (N-terminal to RuvCI, RuvCI-RuvCII, RuvCII-RuvCIII, and RuvCIII to the C-terminus), and multiple sequence alignments were conducted on each with default parameter settings in MUSCLE^24^. A maximum likelihood phylogenetic tree and bootstrap values were derived using Phangorn^25^ and a blocks substitution matrix^26^ (“Blosum62”) model with optimized discrete gamma and proportion variables (Figure 1A).

### Plasmid Construction and Rice Transformation

Cms1-encoding genes were codon optimized for monocot plant codon usage. An N-terminal nuclear localization signal was added to all nuclease genes. The sequence of the monocot optimized version of SmCms1and expression elements can be found in Supplementary Figure 3. Monocot optimized sequences of SuCms1, MiCms1, and ObCms1 can be found in Supplementary Figures 4-6. Plant transformation constructs containing Cms1 gene driven by enhanced 35S promoter, crRNA cassette driven by OsU6 promoter and repair templates containing hygromycin selection cassette, were assembled into three individual vectors, according to previously published methods^19^. Plasmids used in this study can be found in Supplementary Table 4. Biolistic rice (*Oryza sativa* L. cv. Kitaake) transformation was performed according to previously published methods^19^. Briefly, embryogenic rice callus was bombarded with gold particles coated with 2µg of total DNA. The DNA components were loaded at a ratio of 0.5:0.5:1 µg of nuclease:crRNA:repair template. Bombarded rice callus was placed on selection medium containing hygromycin (50mg/L) for 3 weeks in the dark at 32°C. Resistant callus pieces were sub-cultured to fresh media and returned to same conditions for one week prior to sampling and handoff for molecular analysis. Positive events for indel or HDR mutations were placed on regeneration media to generate transgenic events.

### Rice Characterization

DNA extractions from rice leaf and callus tissue, PCR analyses, and T7 Endonuclease I (T7EI) assays were performed as described previously^19^. Sanger sequencing of PCR products amplified from rice callus DNA extracts was performed to validate all putative positive T7EI results.

